# Transposon-insertion sequencing in a clinical isolate of *Legionella pneumophila* identifies essential genes and determinants of natural transformation

**DOI:** 10.1101/2020.10.01.323287

**Authors:** Léo Hardy, Pierre-Alexandre Juan, Bénédicte Coupat-Goutaland, Xavier Charpentier

## Abstract

*Legionella pneumophila* is a Gram-negative bacterium ubiquitous in freshwater environments which, if inhaled, can cause a severe pneumonia in humans. The emergence of *L. pneumophila* is linked to several traits selected in the environment, the acquisition of some of which involved intra- and interkingdown horizontal gene transfer events. Transposon-insertion sequencing (TIS) is a powerful method to identify the genetic basis of selectable traits as well as to identify fitness determinants and essential genes, possible antibiotic targets. TIS has not yet been used to its full power in *L. pneumophila*, possibly because of difficulty to obtain a high-saturation transposon insertion library. Indeed, we found that ST1 isolates, to which belong the commonly used laboratory strains, are poorly permissive to saturating mutagenesis by conjugation-mediated transposon delivery. In contrast, we obtained high-saturation libraries in non-ST1 clinical isolates, offering the prospect of using TIS on unaltered *L. pneumophila* strains. Focusing on one of them, we therefore used TIS to identify essential genes in *L. pneumophila*. We also revealed that TIS could be used to identify genes controlling vertical transmission of mobile genetic elements. We then applied TIS to identify all the genes required for *L. pneumophila* to develop competence and undergo natural transformation, defining the set of major and minor Type IV pilins that are engaged in DNA uptake. This work paves the way for the functional exploration of the *L. pneumophila* genome by TIS and the identification of the genetic basis of other life traits of this species.

**Importance:** *Legionella pneumophila* is the etiologic agent of a severe form of nosocomial and community-acquired pneumonia in humans. *L. pneumophila* is found in man-made and freshwater environments which are the causing source of the infection. The environmental life traits of *L. pneumophila*, such as its abilities to form biofilms, resist biocides and unicellular predators, are essential to its ability to accidentally infect humans. A comprehensive identification of the genetic basis of these life traits could be obtained through the use of transposon-insertion sequencing. Yet, this powerful approach, had not been fully implemented in *L. pneumophila*. Here we described the successful implementation of the transposon-sequencing approach in a clinical isolate of *L. pneumophila*. We identify essential genes, potential drug targets, and genes required for horizontal gene transfer by natural transformation. This work represents an important step towards identifying the genetic basis of the many life traits of this environmental and pathogenic species.

## Introduction

*Legionella pneumophila* is a Gram-negative bacterium, ubiquitous in freshwater environments where it can be found in planktonic form, in biofilm communities or associated to amoebic protozoa which constitute its natural host (1). *L. pneumophila* can resist predation by amoeba and even establish an intracellular vacuole in which it can multiply, while being protected from external environment (2). Man-made water systems have offered a new breeding-ground for the development of *L. pneumophila*. Inhalation by humans of aerosols produced by these systems and contaminated by *L. pneumophila* can cause Legionnaires’ disease (3). This community-acquired disease, which is most often characterized by a severe pneumonia, occurs when *L. pneumophila* infects alveolar macrophages (4). In both macrophages and its natural amoebal hosts, *L. pneumophila* replicates intracellularly by hijacking the host cellular machinery (5). This requires the Icm/Dot type IV system (6, 7), a conjugative system that can secrete up to 300 effector proteins (8, 9). The first genome sequences of the original Philadelphia outbreak strain Philadelphia-1 (10) and the endemic strain Paris (11) provided early evidence of genes encoding eukaryotic-like proteins, some of which are the effector proteins substrates of the Icm/Dot system. Phylogenetic analyses suggest that these genes would have been acquired by inter-kingdom horizontal gene transfer (HGT) during co-evolution of *Legionella* and its natural host for millions of years (12). Hundreds of genome sequences of *L. pneumophila* clinical isolates have now revealed that recombination events are common in this species (13–15). Thus, intra-specific and inter-kingdom HGT events are playing a major role in the evolution and adaptation of this species. The high plasticity of the genomes of *L. pneumophila* is consistent with the fact that it is competent for natural transformation (16). Natural transformation refers to the ability of certain bacteria to capture exogenous DNA and integrate it into their genome by homologous recombination (17). It is one of the driving forces for bacterial evolution that can lead to the emergence of new pathogenic bacteria and new antibiotic-resistant recombinants. It is a widespread mechanism of HGT in bacteria, with more than 80 experimentally-confirmed transformable species (18). The DNA uptake mechanisms and associated proteins constituting the so-called “DNA uptake machinery” are highly conserved (17), suggesting that most species are potentially transformable. DNA uptake first involves a type IV pilus (T4P) (19) whose direct observation supports a model in which it binds DNA via its tip, and its retraction allows the internalization of DNA into the periplasm (20). The periplasmic DNA-binding protein ComEA serves as a ratchet (21, 22) and large amounts of DNA can accumulate in the periplasm before being converted into single-stranded DNA (ssDNA) and translocated across the cytoplasmic membrane through the ComEC inner membrane channel (23). In the cytoplasm, the ssDNA is protected by the transformation-dedicated protein DprA (24) and the single-stranded binding protein SsbB (25). If the internalized ssDNA possesses homologous regions with the bacterial chromosome, it is integrated by homologous recombination mediated by the recombinase RecA which interacts with DprA (26). In Gram-negative bacteria, the newly discovered ComM helicase is also involved in this recombination process (27).

In most transformable species, these proteins are not expressed constitutively but only when the bacterium is in a genetically programmed and transient state called “competence” (28). *L. pneumophila* was first reported competent when grown at 37°C under some form of stress, such under microaerophilic conditions (16) or exposure to DNA-damaging agents (29). In the absence of any stress, *L. pneumophila* becomes transiently competent when grown at 30°C at the transition between the exponential and stationary growth phases (30, 31). *L. pneumophila* is unique in that the regulation of competence does not involve transcriptional activation of the competence regulon. Rather, the core genes encoding the DNA uptake system (*comEC, comEA, comFC, comM*) are subjected to post-transcriptional repression by a ribonucleoprotein complex consisting of a small RNA, RocR, and an RNA chaperone, RocC (31). At the onset of the stationary phase, the expression of RocR decreases and the translation of the mRNAs encoding the DNA uptake system allows *L. pneumophila* to take up and recombine extracellular DNA. Most of *L. pneumophila* clinical isolates do transform under these conditions, yet some isolates fail to develop competence and in some instance, this is due to the presence of a mobile genetic element (MGE) that encodes a RocR homolog that acts as a substitute of the chromosome-encoded RocR (32). Competence is further repressed in stationary phase by the quorum-sensing system (33). The regulation of competence in *L. pneumophila* still remains poorly understood (34).

Regulation of competence is best understood in the Gram-positive *Streptococcus pneumoniae* in which the comprehensive genetic approach of transposon-insertion sequencing has recapitulated decades of findings (35). Beyond the identification of additional regulatory or functional elements of natural transformation, such approach allowed for a better understanding of the biology of this bacterium by identifying genes involved in virulence and in resistance against stresses. Transposon-insertion sequencing (TIS) approaches encompass a number of similar methods (Tn-seq, TraDIS, INseq, HITS) (36–39) that have been used for the identification of essential genes on a genome-wide scale in a number of species (40). TIS relies on the mapping and quantification of transposon insertion mutants by high-throughput DNA sequencing and a critical factor is to obtain high-saturation libraries of transposition mutants (41). TIS has recently been applied to *L. pneumophila* with a focus on effector-encoding genes and their conditional involvement in intracellular replication (42, 43). However, the libraries of mutants were either targeted for effectors (42) or of low coverage (43). Thus, the full power of TIS has not yet been harnessed to understand fundamental or specific aspects of the biology of *L. pneumophila*, possibly because of the difficulty of obtaining high-saturation mutant libraries. In addition, the current libraries were constructed in the lp02 strain which has lost competence regulation during its laboratory domestication (44). Here we sought to obtain a high coverage library for Tn-seq in *L. pneumophila* that could be used to apprehend the genetic basis the many life traits of this species. We found that some clinical isolates of *L. pneumophila* are more permissive to transposon mutagenesis than the commonly used laboratory strains. We obtained a high coverage Tn-seq library in an unaltered clinical isolate, and identified genes essential for fitness and growth in axenic medium. We then applied Tn-seq to identify the genes involved in competence and natural transformation.

## Results and discussion

### A high-saturation Tn-seq library of L. pneumophila

With the objective of obtaining a Tn-seq library of *L. pneumophila*, we tested the conjugative delivery of the Himar1-based transposon encoded by the *pir*-dependent mobilizable plasmid pBT20 to the commonly used strain Paris. Conjugation assays with the MFDpir donor strain only produced a handful of insertional mutants. We hypothesized that the Paris strain was particularly resistant and tested 12 other clinical isolates belonging to the sequence type (ST) 1. Similarly to the Paris strain, none of the ST1 isolates generated a meaningful number of mutants. We concluded that for an unknown reason the ST1 isolates (which would include the Philadelphia-1 derived laboratory strains lp02 and JR32) were poorly permissive to conjugative transfer and/or to transposition by Himar1. We thus tested 8 other non-ST1 clinical isolates. We obtained several thousands mutants for 5 of these. We decided to purse with isolate HL-0709-3014, for which we obtained a complete genome composed of a circular chromosome of 3,405 kb and a plasmid of 106 kb (see Material and Methods). 3,183 open-reading frames were detected, 2,791 and 2,741 of which have orthologs in the Paris and Philadelphia-1 strains, respectively (Dataset S1). HL-0709-3014 belongs to the ST18 lineage, which is closely related to the ST1 lineage. Hence, it is phenotypically similar to the Paris strain, it is naturally transformable and shows similar intracellular replication rates in amoebae (Fig. S1). It also effectively replicates in human and murine macrophages (>2 log growth in 72 h) (Fig. S1). We isolated HL77S, a spontaneous streptomycin-resistant mutant of HL-0709-3014, and subjected it to mutagenesis with the transposon of pBT20. This mariner-based transposon inserts at TA sites and includes an outward facing Ptac promoter that can minimize possible polar effects on operon and downstream genes. About 250,000 colonies of mutants were isolated on CYE plates and collected (initial isolation). The library was then cultured in rich medium at 30°C and re-isolated on CYE (second isolation). Sequencing of the transposon insertion sites revealed a maximum of 110,679 unique insertion out of 255,021 possible TA sites (43% saturation) and an average of one insertion site every 31 bp. This represents a significant improvement over the previously published library in the ST1 lp02 strain which consisted in 17,781 unique insertions sites (7% saturation) (43). Thus, we obtained a high saturation Tn-seq library in an *L. pneumophila* clinical isolate that can be used as a surrogate to the commonly used laboratory strains (Paris, JR32, lp02, AA100).

### Analysis of gene essentiality

The high saturation allowed the identification of genes essential for growth. To do so, we used two statistical methods; the Gumbel method (45), a Bayesian model based on longest consecutive sequence of TA sites without insertion in the genes, and the HMM method based on the detection of genes with unusually low read counts (46). Both methods gave similar results with 401 (Gumbel) and 500 genes (HMM) identified as essential, 382 of which were identified as essential by both methods (Dataset S1). This is consistent with the average number (391) of essential genes identified in other bacterial species (47). The data confirmed our previous observation of the essentiality of the actin-like protein MreB (48) but also of MreC and MreD, while intergenic insertions between *mreC* and *mreD* are tolerated (Fig. 1A). Comparative analysis of the second and initial isolation, identified 181 genes non-essential at the initial isolation but whose inactivation impaired fitness (log2FC<-2, P<0.05). These include the gene encoding the exoribonuclease R, whose growth defect was previously reported (49), the RNA chaperone Hfq, but also more surprisingly the substrates of the Icm/Dot Type IV secretion AnkQ and SdbB (Dataset S1). Presumably because their inactivation lowered the fitness so dramatically, 61 genes not essential after the initial isolation were deemed essential on the second isolation. For instance, these include the phosphoenolpyruvate synthase-endoding gene *ppsa* (Fig. 1A), the genes encoding the sigma factor RpoS, the RecA recombinase, the tyrosine recombinase XerC involved in chromosome dimers resolution and the tmRNA-binding protein SmpB involved in trans-translation. Indeed, this is consistent with our previous demonstration that trans-translation is essential in *L. pneumophila* (50) and that no insertions are observed in the tmRNA-encoding gene (Fig. 1B). The vast majority of the essential genes have orthologs in the Paris and Philadelphia-1 genome, as expected for genes that encodes proteins involved in the fundamental processes of the cell. However, of all genes found to be essential either on initial or second isolation, 14 have no orthologs in the Paris and Philadelphia-1 strains. How could strain-specific genes be essential? Three of these genes (HL77S_01135, HL77S_01141, HL77S_01146) encode antitoxin component of toxin-antitoxin (TA) modules clustered within a 4 kb segment. HL77S_01068 is located next to a gene encoding a toxin of type II TA system, suggesting that it also encodes an antitoxin. Others have no known function, such as HL77S_02141 which encodes a protein with a predicted helix-turn-helix (HTH) motif or HL77S_00079 encoding a protein with a conserved domain of unknown function (DUF3800). Consistently with being part of the accessory genome, genomic comparison with other complete genome of *L. pneumophila* indicate that all of these genes reside in highly variable regions, often in proximity to putative transposase and prophage integrase. Yet, no genetic structure corresponding to a complete mobile genetic element (MGE) could be detected. In contrast, two essential genes (HL77S_00162, HL77S_00189) are within a recognizable MGE corresponding to an integrative conjugative element (ICE) inserted at the 3’ end of the tmRNA-encoding gene and carrying a conjugative system homologous to the Lvh system (Fig. 1B). HL77S_00162 is an HTH-type regulator and HL77S_00189 is predicted to encode a DNA-cytosine methyltransferase. The ICE shows a third essential gene (HL77S_00187, *prpA*), conserved in the lvh ICE of the Paris strain and encoding a LexA/CI-like repressor homolog (Fig. 1B). Another strain-specific essential gene (HL77S_00197) is located just downstream of the ICE, in a unique region that may represent a remnant of another MGE. Another three of the 14 strain-specific essential genes (HL77S_03181, HL77S_03182, HL77S_03183) are part of the 106-kb conjugative plasmid and clustered with another essential gene (HL77S_03071) which has a homolog on the pLPP plasmid of the Paris strain (Fig. 1C). Encoding Rep or Par homologs, these genes are involved in replication/partition of the plasmid and their inactivation likely resulted in plasmid loss. On this plasmid, two other genes of unknown function also appear essential, one that is unique and with no conserved domain (HL77S_03140) and one with a homolog on pLPP (HL77S_03116, *plpp0094*) containing an N-terminal HTH motif and a C-terminal nucleotidyltransferase (NT) domain also found in DNA polymerase beta (Fig. 1C). Insertions in two divergently oriented genes (HL77S_03176, HL77S_03177) are also associated with strong fitness defects and are of unknown function (Fig. 1C). Overall, we found that many essential genes can be found within MGEs. Insertion in genes controlling vertical transmission can result in the loss of the MGE (and thus of transposon-insertions making the corresponding gene seemingly essential). This might be the case for repressor of excision of ICE or genes required for replication/partition of plasmids. Other genes might be required to limit the cost of the MGE on the fitness of their host. This might be the case for the LexA/CI-like repressor of ICE, as exemplified by the *Vibrio cholerae* SXT ICE, for which inactivation of the LexA/CI-like repressor SetR is deleterious to its host (51). Whatever the mechanism, the genes characterized as essential in MGEs ensure their vertical transmission. Thus, our result indicates that in addition to identifying genes required for the fundamental functions of the cell, Tn-seq analyses can also reveal novel genes that contribute to vertical transmission of MGEs, representing an untapped resource to study the bacteria-MGE co-evolution.

**Figure 1.**
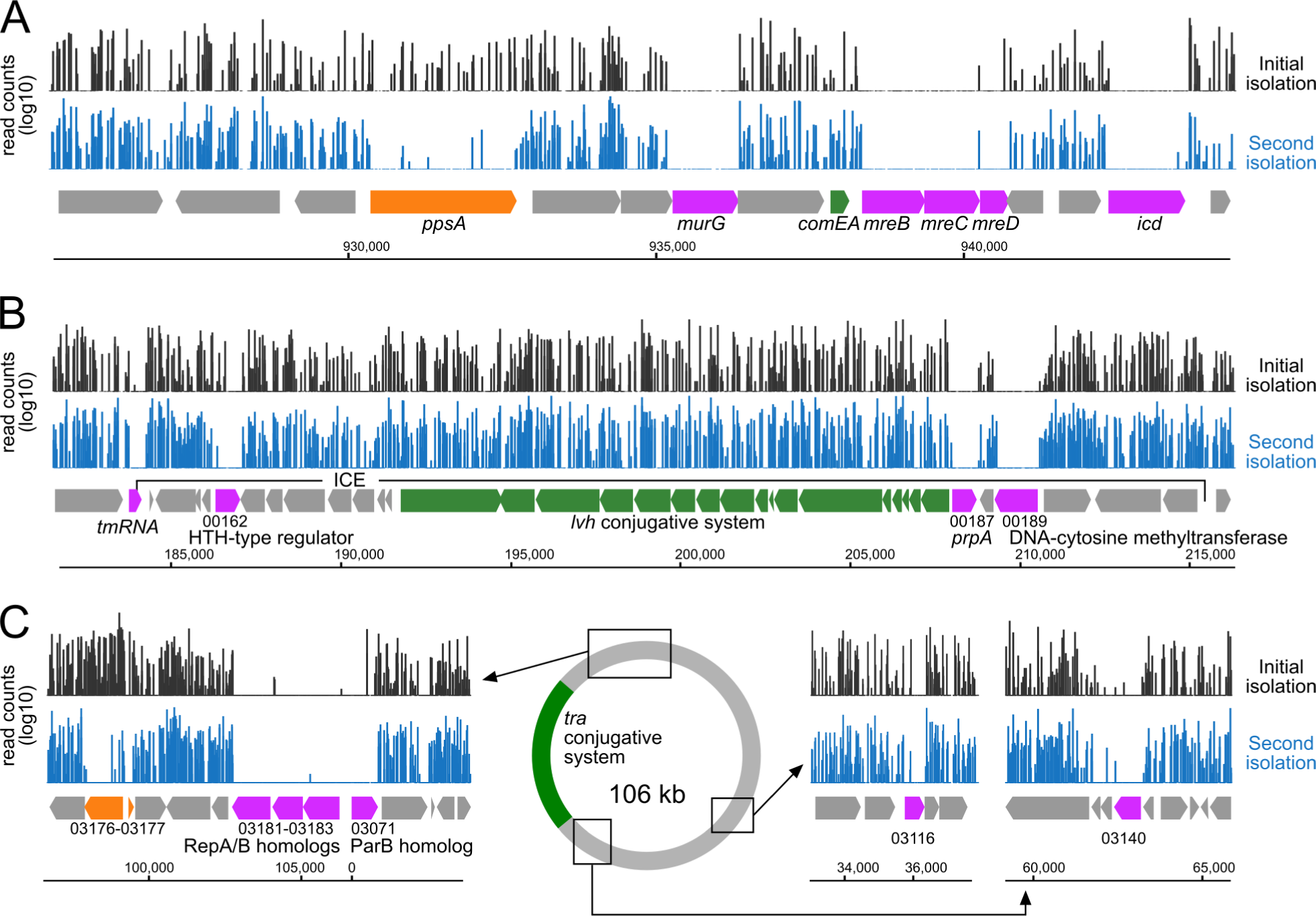
Tn-seq analysis of *L. pneumophila* strain HL77S. A) Log10 reads counts of transposon insertions after initial library isolation (black) and second isolation (blue). Genes identified as essential are colored in magenta, fitness determinants are colored in orange. Other genes of interest are colored in green. B) Transposon-insertion coverage in a region encompassing an integrative conjugative element (ICE) harboring essential genes (magenta) and genes encoding a conjugative system (green). The duplicated sequence gcgggttcgattcccgccgcctccacca of the tmRNA and located 66 kb away delineate the boundaries of the ICE. C) Essential genes and fitness determinants in the conjugative plasmid of HL77S.

### Tn-seq analysis of natural transformation in L. pneumophila

We seek to use the Tn-seq library to identify all genes required for competence and subsequent natural transformation. Mutants defective for expression of competence, DNA uptake, protection or recombination would not be able to undergo transformation and would thus be missing in the transformed population. We subjected the Tn-seq library to natural transformation with two distinct transforming DNA carrying a kanamycin resistance cassette inserted in the *legk2* gene (encoding an Icm/Dot substrate) or in the *ihfB* gene (encoding the B subunit of the integration host factor IHF). This strategy should limit false positive arising from epistatic interactions between the gene in which the selected resistance cassette is inserted (*legk2* or *ihfB*) and the transposon-disrupted genes. Transformation frequencies of the HL77S Tn-seq library were in the range of 1×10^−5^-6×10^−4^ and to avoid a bottleneck effect (41) we collected over 5×10^6^ transformants. The control, non-transformed populations were sub-sampled to obtain a similar number of isolated colonies. We observed 28 genes in which insertions cause a decrease (log2FC<2, P<0.05) of the mutants in the population transformed with either the *legK2*::kan or the *ihfB*::kan DNA (Fig. 2A). As expected, these include the gene encoding the periplasmic DNA receptor ComEA, the genes required for DNA transport across the inner membrane (*comEC, comFC*), for ssDNA protection in the cytoplasm (*dprA*) and recombination (*comM*). Many of the other genes encode factors known to be involved in Type IV pilus assembly, confirming the role of this system in natural transformation of *L. pneumophila* (16). These include the retraction ATPase PilT (lpp1995), the extension ATPase PilB, the PilQ secretin, the PilC platform protein and the proteins of the PilMNOP complex. Interestingly, we observed no transformation defect for insertions in the gene encoding another putative pilus retraction ATPase (lpp2271). Two putative pilins, pilE (lpp0681) and pilA_2 (lpp1890) were also identified as essential for transformation. Other genes potentially involved in natural transformation or regulation of competence were also identified (*lpp1976, lpp1977, lpp1978, lpp3030, lpp2632, djlA, letA, letS*). In order to confirm their role and disentangle their involvement in DNA uptake or in regulation, we constructed gene deletion mutants in a Paris strain with a premature stop codon of the RocC chaperone (Paris *rocC*_*TAA*_) which is defective for repression of competence and constitutively transformable (31). Deletion mutants corresponding to genes known to be involved in natural transformation were defective for transformation as expected (Fig. 2B). The *comEC* and *comFC* mutants were totally defective for transformation, the *comM* and *comEA* mutants showed a ∼100-fold decrease in transformation frequencies as observed for other species (27, 52). Similar partial transformation defects were observed for mutants of *djlA*, encoding a DnaJ-like protein required for intracellular replication in *Legionella dumoffii (53), a*nd *lpp3030*, a *Legionellaceae*-specific gene encoding an uncharacterized protein with a putative signal peptide. However, in this constitutively competent background we could not confirm the involvement of *lpp2632* which encodes a glutaryl-CoA dehydrogenase, indicating that this gene is dispensable for the transformation process (Fig. 2B). Mutants of this gene show a reduced fitness (log2FC=-1.99, P<0.01) (Dataset S1), suggesting that the transformation defect observed in the Tn-seq analysis is an indirect consequence of the mutants limited growth that could prevent entry in the competence state at the onset of the stationary phase. Intriguingly, in this constitutively competent strain, a deletion mutant of *letS* also showed no transformation defect (Fig. 2B). LetS is the sensor of the LetA/LetS two-component system (TCS) homologous to the BarA/ UvrY system in *Escherichia coli* (54) and GacS/GacA in *Pseudomonas* spp (55). In *L. pneumophila*, the LetA/ LetS system has been identified for the first time in a screen of mutants deficient in the expression of flagellin (56) and has since been shown to be involved in the activation of various virulence traits as well as intracellular growth in amoeba (57–60). One of the major roles of the LetA/S TCS is to enable the transition from the transmissive to the replicative phase (61). The facts that both LetA and LetS output together in the transformation screens while the *rocC*_TAA_ Δ*letS* mutant is not defective for transformation suggests that this TCS is involved in the regulation of competence in *L. pneumophila*. To test this, we reconstructed an insertion mutant of the *letA* gene encoding the activator of this TCS in the Paris strain and *rocC*_*TAA*_ genetic backgrounds and tested them for their ability to undergo transformation. Consistent with the Tn-seq data, inactivation of LetA in the Paris strain reduced transformability by over 500-fold (Fig. 2C). In contrast, like the Δ*letS* mutant, the Δ*letA* mutant in the constitutively competent strain *rocC*_TAA_ is only marginally affected for natural transformation (Fig. 2C). These data suggest that the LetA/S TCS is involved in the regulation of competence upstream of the regulation controlled by the RocC/RocR system. Further work will be needed to determine the precise role of this TCS and the associated regulatory cascades in the regulation of *L. pneumophila* competence.

**Figure 2.**
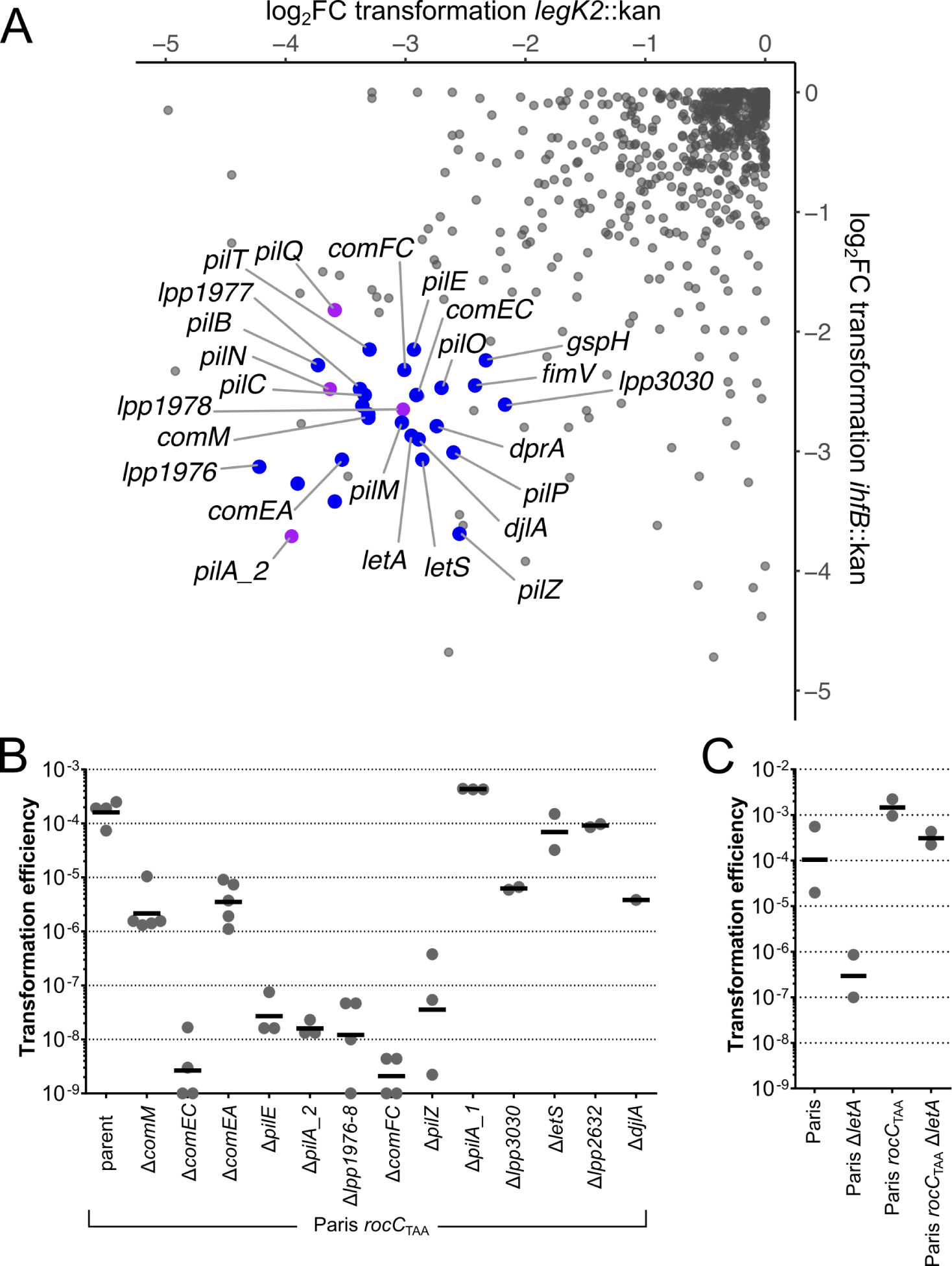
Identification of genes required for natural transformation by Tn-seq. A) Scatter plot of fold-change (log_2_) of insertions in the corresponding genes in two tested transformation conditions. HL77S was subjected to natural transformation with a 4kb-PCR fragment of the *legK2* or *ihfB* genes interrupted by a kanamycin resistance gene. Reads count per genes were determined and expressed as fold-change between the non-transformed population and the *legK2*::*kan* or *ihfB*::*kan* transformed populations. Individual genes (gray dots) were considered required for natural transformation if log_2_FC was >2 or <-2 and if P<0.05 in one (magenta dots) or both conditions (blue dots). B) Natural transformation efficiency of reconstructed mutants in the Paris *rocC*_TAA_ strain which is constitutively competent for natural transformation. Transformation experiments were performed at least three times independently and transformation frequencies were plotted (grey dots) along with the geometric mean (black line). C) Natural transformation efficiency of the reconstructed mutant Δ *letA* in the original Paris strain and constitutively competent Paris *rocC*_TAA_ strain. Transformation experiments were performed twice independently and transformation frequencies were plotted (grey dots) along with the geometric mean (black line).

### Major and minor pilins required for natural transformation

With the remarkable exception of *Helicobacter pylori* (62), in all Gram-negative bacteria DNA uptake requires type IV pili (19). Type IV pili are extracellular filaments resulting from the assembly of thousands copies of an abundant major pilin but also of less abundant minor pilins that could be embedded in the filaments (core minor pilins) or at its tip (non-core minor pilins) (63, 64). The nomenclature of pilins is relatively confusing but the major pilin is generally called PilA, although in *Neisseria sp*. that protein is called PilE (64). In addition, some species carry multiple copy of pilins and at least in *Thermus thermophilus* two major pilins (PilA4 and PilA5) are assembled into distinct filaments respectively required for natural transformation and twitching motility (65). The *L. pneumophila* genomes show two putative PilA homologs encoded by two consecutive genes (*pilA_1*, lpp1889; *pilA_2*, lpp1890) in a locus away from any other genes encoding Type IV pilus components. Both the Tn-seq data and reconstructed mutants show that PilA_2 is required for natural transformation while PilA_1 is dispensable (Fig. 2A and 2B). The two copies of PilA-encoding genes may have resulted from a gene duplication event, followed by the loss of function of one of the two copies. The Tn-seq data show that a putative pilin PilE (Lpp0681) appears required for transformation, while five genes upstream of pilE (*lpp0686*-*lpp0682*) and respectively annotated as PilC/PilY1 and minor pilins PilX, PilW, PilV and GspH/FimT appears dispensable. Targeted gene deletion also confirmed the Tn-seq result that PilE is required for natural transformation, corroborating an initial observation that a mutant of the *pilE* gene (then denoted *pilE*_*L*_) is not competent for transformation (16). Based on sequence comparison with PilA from *P. aeruginosa, pilE*_*L*_ was then proposed to encode a type IV pilin structural gene (66). We thus investigated which of PilE and PilA_2 constitute the major pilin in *L. pneumophila*. We tested the complementation of the Δ*pilE* and Δ*pilA_2* deletion mutants obtained in the constitutively competent strain *rocC*_TAA_. Both *pilE* (*lpp0681*) and *pilA_2* (*lpp1890*) were ectopically expressed from an IPTG-inducible promoter to produce fusion proteins with a C-terminal FLAG epitope. Western-blot analysis showed that both PilE-FLAG and PilA_2-FLAG could be expressed in an IPTG-dependent manner with PilE-FLAG always expressed at a higher level than PilA_2-FLAG, likely reflecting the efficiency of their ribosome-binding site (Fig. 3A). Data show that a low expression of PilE-FLAG is sufficient to restore natural transformation in the *rocC*_TAA_ Δ*pilE* mutant as full complementation of the transformation phenotype is obtained even in the absence of IPTG (Fig. 3B). In contrast, a higher concentration of IPTG and thus a higher expression of PilA_2-FLAG is required to obtain a functional complementation of the *rocC*_TAA_ Δ*pilA_2* mutant (Fig. 3B). The results are consistent with a model in which PilA_2 is the major pilin while PilE is a low-abundance minor pilin. In addition, when expressed ectopically in the *rocC*_TAA_ strain, PilA_2-FLAG assembles in long extracellular filaments (Fig. 4A and 4B). In the *rocC*_TAA_ Δ*pilE* strain, fewer PilA_2-FLAG filaments are observed by microscopy and western-blot confirmed a lower abundance of extracellular PilA_2-FLAG (Fig. 4A and 4B). This indicates that PilE, while not strictly essential still plays a role in pilus formation. Minor pilins have been proposed to localized at the tip of the pilus and stabilize it (63). In *Vibrio cholerae*, DNA binding has been observed to occur at the tip of the pilus (20). Because PilE is not strictly essential for pilus assembly but required for transformation and DNA internalisation (Fig. 3B and 4C), we propose that PilE is the DNA receptor at the tip of a pilus composed of PilA_2 subunits.

**Figure 3.**
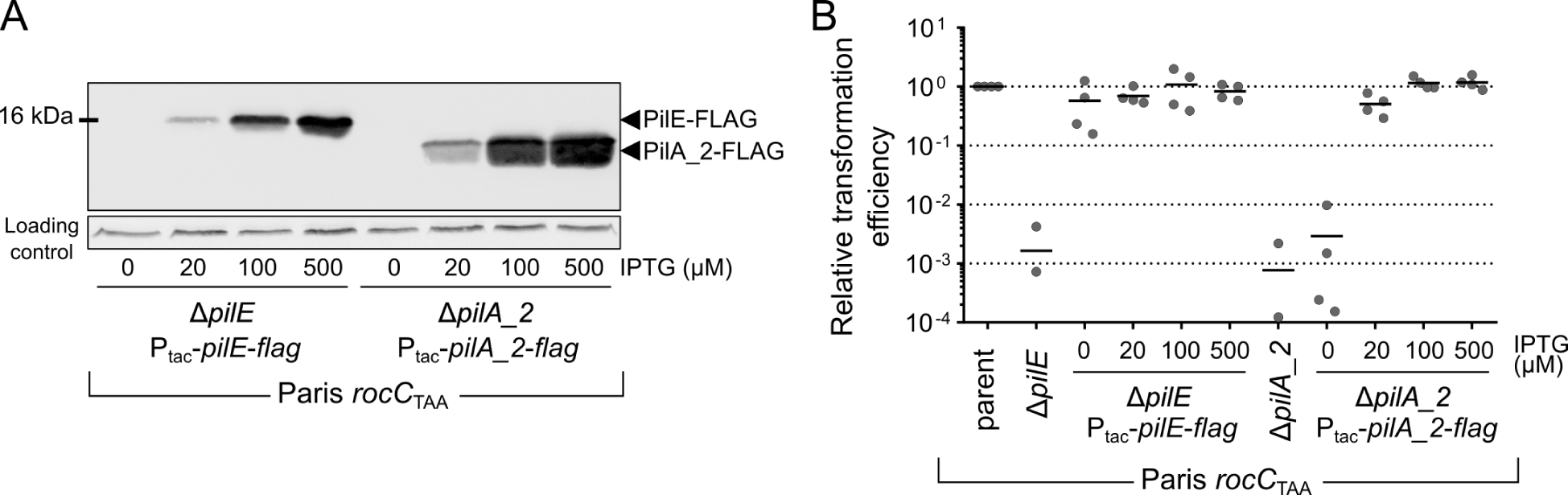
PilA_2 is the major pilin of the *L. pneumophila* transformation pilus. A) Western-blot analysis of ectopically-expressed PilA_2-FLAG (encoded by p1890F) and PilE (encoded by p0681F) as a function of the IPTG inducer. B) Complementation of the Δ*pilE* and Δ*pilA_2* mutants in the Paris *rocC*_TAA_ strain by the ectopic expression of PilA_2-FLAG (encoded by p1890F) and PilE (encoded by p0681F). Transformation frequencies were determined four times independently as a function of the IPTG inducer, and normalized to 1 for the parental strain (Paris *rocC*_TAA_).

**Figure 4.**
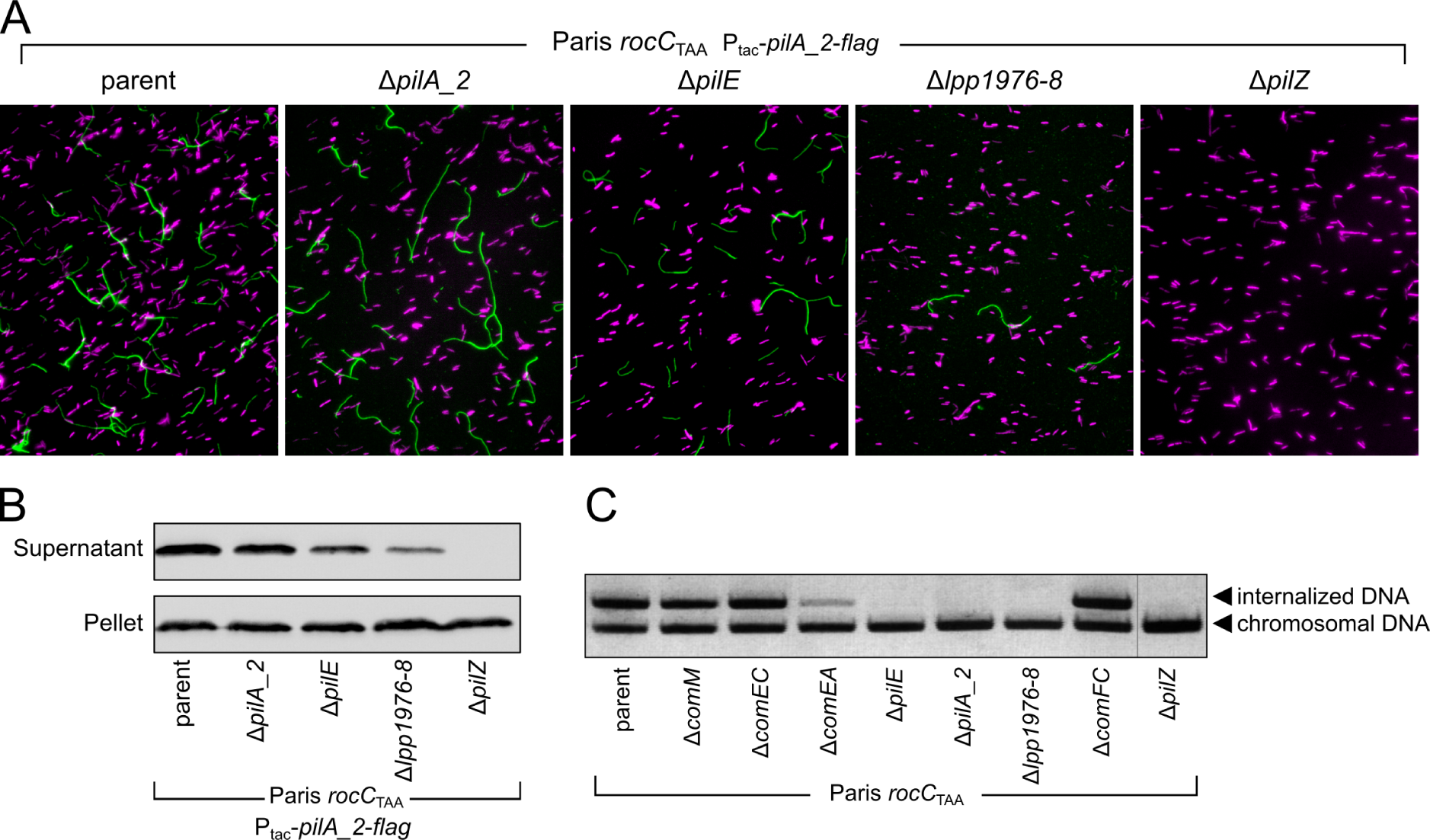
PilA_2 assembly into extracellular filaments depends on *pilE*, the operon *lpp1976-8* and *pilZ*. A) Visualization of PilA_2-FLAG filaments (green) by immunofluorescence microscopy using fluorescein-conjugated anti-FLAG antibody. Bacteria were visualized by labeling DNA with Hoechst 33288 (magenta). B) Western-blot detection of extracellular PilA_2. Bacteria were vortexed to release pili which were precipitated from supernatants. PilA_2-FLAG was detected in supernatant and whole cell lysates (pellet) using Anti-FLAG antibodies. C) DNA uptake assay of the reconstructed mutants defective for natural transformation. The ability of the transformation-deficient mutants were tested for the ability to internalize pGEM-HYG1, a non-replicative plasmid. Following incubation with the DNA and subsequent DNAse I treatment, the internalized DNA was detected in cells by PCR for pGEM-HYG. As control, chromosomal DNA was also detected by PCR. This multiplex PCR was analyzed by agarose gel electrophoresis and labeling of DNA with ethidium bromide.

### Genes of unknown function and pilZ

In addition to the pilin mutants that were strongly defective for natural transformation, we investigated the underlying reason for the strong transformation defect of the mutant deleted of the operon *lpp1976*-*lpp1977*-*lpp1978* (Fig. 2B). Consistent with being important for natural transformation this operon was found to be up-regulated in the constitutively transformable mutant *rocC*_TAA_ (31). Automated annotation did not assign a predicted function for the three genes and blast search failed to identify homologs outside of the *Legionella* genus. The deletion mutant of the entire operon Δ*lpp1976-8* was found unable to take up DNA (Fig. 4C), indicative of a defect in Type IV pilus-mediated DNA import. Indeed, in this mutant, the levels of extracellular PilA_2-FLAG were strongly reduced (Fig. 4B). The mutant produced few, and short, PilA_2-FLAG filaments (Fig. 4A) revealing a major defect in Type IV pilus assembly or stability. A search for conserved domains in the three predicted proteins only identified, in the 268 aa-long Lpp1977, a partial homology with the N-terminal part of the Tfp pilus assembly protein PilW. This suggested that *lpp1976*-*lpp1977*-*lpp1978* would encode a set of minor pilins. Indeed, PilFind (67) identified an N-terminal transmembrane segment in all three predicted proteins and a type III signal in Lpp1977 and Lpp1978. The operonic organization of these three genes is reminiscent of the operon encoding four minor pilins of the type IV pilus of *N. meningitidis* (*pilHIJK*) and *P. aeruginosa* (*fimU-pilVWX*) and of the Type II secretion system (T2SS) of enterotoxigenic *E. coli* (*gspHIJK*). In the latter system, the last three genes (*gspIJK*) encode minor pseudopilins which assemble into a stable complex (68). This complex of minor pilins would form in the inner membrane to establish a platform for the assembly of the major pilin (69), and remain at the tip of the pilus, stabilizing it (63). Such heterotrimeric complex may be formed by minor pilins of limited homology but displaying structural similarity (70). Altogether, this supports the hypothesis that in *Legionella* species, the initiation complex of the transformation pilus is formed by Lpp1976, Lpp1977 and Lpp1978 which serve as a scaffold for assembly of the major pilin PilA_2.

Another gene whose deletion resulted in strong deficiency in natural transformation is *pilZ*. The Δ*pilZ* mutant is defective for DNA uptake and is totally unable to produce extracellular PilA_2 or assemble PilA_2 filaments (Fig. 4). PilZ was originally identified in *P. aeruginosa* as required for the secretion of PilA polymers, pilus genesis and Type IV pilus-dependent motility (71). Although the *P. aeruginosa* PilZ served as the founding member of a diverse family of proteins with PilZ domains (72), some of which bind the cyclic-di-GMP second messenger, it itself does not bind c-di-GMP (73). A *pilZ* mutant in *Xanthomonas campestris* pv. campestris displays a minor defect in Type IV pilus-dependent motility (74) and this PilZ ortholog directly binds to the PilB ATPase and the c-di-GMP interacting FimX protein (75). Yet, no homolog for FimX could be identified in *L. pneumophila* and Tn-seq did not reveal any c-di-GMP synthesis enzyme required for natural transformation. However, Tn-seq did show that PilB was important for transformation (Fig. 2A). We thus speculate that, in *L. pneumophila*, PilZ controls Type IV pilus assembly independently of c-di-GMP signaling and through a direct interaction with PilB.

## Conclusion

We report on a clinical isolate of *L. pneumophila*, which displays phenotypes (intracellular replication, competence for natural transformation) similar to commonly used laboratory strains. In contrast to laboratory strains, a high-saturation Tn-seq library could be obtained and allowed to define essential genes, including strain-specific genes in MGEs. Tn-seq analyses of transformation, with follow-up work performed in the Paris strain, defined the set of major and minor Type IV pilins that are engaged in DNA uptake. While we here focused on mutants that were strongly deficient for natural transformation, Tn-seq also identified potential regulators of competence as well as genes of unknown function that also participate in natural transformation (for instance *djlA* and *lpp3030*). We exemplify here that strain HL77S could represent a surrogate for the commonly used lab strains to perform Tn-seq analysis. Unleashing the full power of Tn-seq is a major step toward the identification of the genetic basis of traits that turned *L. pneumophila* into a successful pathogen, such as its ability to form biofilms, resist biocides and unicellular predators.

## Material and methods

### Bacterial strains and growth conditions

*Legionella pneumophila* strains were grown in liquid medium ACES [N-(2-acetamido)-2-aminoethanesulfonic acid]-buffered yeast extract (AYE) or on solid medium ACES-buffered charcoal yeast extract (CYE) plates. When appropriate kanamycin, gentamycin and streptomycin were added respectively at 15 µg/mL, 10 µg/mL and 50 µg/mL. Clinical isolates of *L. pneumophila*, including HL-0709-3014 (“HL77”), were provided by the Centre National de Référence des Légionelles, Lyon, France. A streptomycin-resistant of HL-0709-3014 mutant was obtained by plating 1 mL of culture on streptomycin-containing CYE plate and named HL77S. The *Escherichia coli* MFDpir (76) with chromosome-integrated RP4 conjugative system was used as donor strain for conjugative transfer of the mutagenesis system pBT20 (77) that carries a Himar1 transposon bearing a gentamycin resistance gene and an outward-facing promoter. MFDpir is auxotrophic for diaminopimelic acid (DAP) and thus was always cultivated with 1% DAP. Axenic *Acanthamoeba castellanii* cells were grown in PYG medium (Proteose yeast extract glucose medium) at 30°C and split once or twice a week. Human U937 cells were maintained in RPMI 1640 with 10 % heat-inactivated fetal calf serum and 1 % penicillin/streptomycin at 37°C and 5 % CO2. Differentiation into macrophages was induced by the addition of PMA (phorbol 12-myristate 13-acetate) at a final concentration of 100 ng/ml. Murine macrophages RAW 264.7 were cultured in DMEM medium supplemented with 10% FBS and 1% penicillin/streptomycin at 37°C and 5 % CO2.

### Intracellular growth experiments

The ability of HL77S strain to infect host cells was compared to the Paris strain. Paris Δ*dotA* was used as a negative control as this mutant is unable to multiply in host cells. The ability of HL77S to replicate in the amoeba *Acanthamoeba castellanii* was determined as follows. Amoebas were resuspended in PYG medium at a concentration of 1.10^6^ amoebas/mL. The suspension was distributed in a flat-bottom 6-well plate (2 mL per well, 2.10^6^ amoebas per well) and incubated 3 h at 30°C to allow amoebas to settle and adhere to the plate. 1 mL of a PY medium suspension containing 2.10^6^ bacteria (from a culture in stationary phase OD∼5) were added in each well to obtain the multiplicity of infection (MOI) of 1. The plate was centrifuged 10 min at 650 g and incubated at 30°C for 72 h. At T=0, 48 h and 72 h, 250 µL of supernatant of each well were serial-diluted and spotted onto CYE plates and incubated at 37°C for 72 h to determine the number of colony-forming units (CFU) per mL. The ability of *L. pneumophila* strains to infect macrophages was determined as follows. Overnight cultures of bacterial strains (OD∼5 in AYE medium) were diluted (1:10) in the appropriate cell culture media (DMEM for RAW 264.7 and RPMI 1640 for U937) and incubated for 1 h at 37°C. Host cells (differentiated U937 and RAW 264.7) were seeded in 24-well plates, 3 wells per condition. Cells were washed and were infected with bacteria at a MOI of 1 or 10. Plates were centrifuged at 500 g for 5 min to promote bacteria-cell contact and incubated at 37°C and 5% CO2 for 24 h to 72 h. Every 24 h, the content of one well per condition was transferred to an 1.5 mL tube and centrifuged at 16,000 g for 5 min. The pellet containing the infected macrophages was resuspended in sterile distilled water to lyse the macrophages and release the bacteria. The suspension was serial-diluted and spotted onto CYE plates and incubated at 37°C for 72 h to determine the titer in CFU/mL.

### Plasmid and strains constructions

Plasmid pJET1.2-legk2::kan, used for natural transformation experiments, was constructed by cloning a 6 kb-long fragment consisting of the *legk2::kan* gene (78) and 2 kb of its flanking regions in pJET1.2/blunt cloning vector (ThermoFisher) according to the manufacturer’s instructions. All the mutants generated in this study are derived from *L. pneumophila* Paris or *L. pneumophila* Paris *rocC*_TAA_. All the genes suspected to be involved in natural transformation were deleted by replacement with a kanamycin resistance gene. To do so, the upstream (PCRA, 2 kB) and downstream (PCRC, 2 kB) regions of each suspected genes were amplified respectively with the primers pairs X_P1/X_P2-tail-pKD4 and X_P3-tail-pKD4/X_P4 (where X designated the genes to be deleted). X_P2-tail-pKD4 and X_P3-tail-pKD4 carrying 30-nucleotide sequences complementary to the ends of the kanamycin cassette. This complementarity was used to assemble PCRA and PCRC to the kanamycin resistance cassette (PCRB, 1,490 kB amplified from plasmid pGEMPKD4 (31) with primers pair pKD4_P1/pKD4_P2) by overlap extension PCR. Overlapping PCRs were naturally transformed in the desired strain. Transformants were selected on CYE supplemented with kanamycin (15 µg/mL). Integration of the kan cassette at the correct locus was finally verified by colony PCR. Plasmids p1890F and p0681F, encoding the FLAG-tagged PilA_2 and PilE, were constructed by amplifying *lpp1890* (pilA_2) and *lpp0681* (pilE) with primers lpp1890-F/lpp1890F-R and lpp0681-F/lpp0681F-R, respectively. The PCR products and the recipient plasmid pMMB207C were digested with *Hin*dIII/*Bam*HI and ligated to place the genes under the Ptac promoter. All strains, plasmids and oligonucleotides are listed in Table S1.

### Generation of transposon insertion mutants library of Legionella pneumophila

Transposon-based random mutagenesis was performed as previously described (79) by conjugative delivery of the Himar1-based transposon suicide vector pBT20 from the donor strain *E. coli* MFDpir to the recipient strain of *L. pneumophila* to be mutagenized. To do so, both bacteria were cultivated overnight at 37°C with shaking in their corresponding liquid media: 7.5 mL LB broth containing 100 µg/mL ampicillin and 1% DAP for *E. coli* and 15 mL standard AYE medium for *L. pneumophila*. Once in stationary phase (i.e; DO∼5), the *L. pneumophila* and *E. coli* cultures were concentrated by centrifugation (5,000 g, 10 min) and cell pellets were resuspended respectively in 1.5 mL sterile water and 0.750 mL sterile PBS. To promote cell-to-cell contacts and the subsequent conjugation, both concentrated cultures were mixed together by pipetting, and spotted on CYE plates without iron and cystein but supplemented with DAP (CYED) (79) until the sample was exhausted. Plates were incubated at 37°C for 5 to 6 h. All the spots were resuspended in sterile water and used to inoculate transconjugants-selective plates (*i*.*e*., CYE plates supplemented with 10 µg/mL of gentamicin). In parallel, the suspension was ten-fold diluted and spotted onto transconjugants-selective plates to evaluate the number of mutants in the library. After 72 h of incubation at 37°C, mutant library was obtained by collecting all colonies from the plates and resuspending them in AYE-15% glycerol. The suspension was aliquoted and stored at −80°C until its use for a Tn-seq screen. This library, called “initial isolation” is named sample XRCR13.

### Natural transformation Tn-seq screen

Transposon mutants of *L. pneumophila* HL77S were screened for their ability to undergo transformation. To avoid a bottleneck effect, a volume of the −80°C frozen library containing ten times the number of mutants in the library was spotted on a CYE plate supplemented with gentamycin (10 µg/mL) and streptomycin (50 µg/ mL) to obtain exponentially growing cells. After 24 h of incubation at 37°C, fresh bacteria from the spot were resuspended in AYE to an OD∼0.2. This suspension was used to perform transformation assays using 2 µg/ mL of either pGEM-ihfB-kan or pGET1.2-legK2-kan as transforming DNA, both conferring resistance to kanamycin. For both transforming DNA, the transformation screen was conducted in duplicate. The suspensions were cultivated at 30°C with shaking for 40 h to ensure that bacteria undergo transformation and achieve an OD∼5. These conditions were expected to give about ∼10^5^ transformants/mL for the “DNA” conditions. Regarding the “no DNA” condition, cultures were diluted in AYE to obtain the same number of CFUs on non-selective plates than the output condition on selective plates. Each sample (“+DNA” and “no DNA”) was used to inoculate respectively nonselective (*i*.*e*., CYE) and transformants-selective (*i*.*e*., CYE supplemented with 20 µg/mL of kanamycin) plates. In parallel, “no DNA” samples were tenfold diluted and spotted on nonselective plates to determine transformation frequencies as mentioned above. After 72 h of incubation at 37°C, colonies were collected from the plates and resuspended in AYE-15% glycerol until the preparation of DNA libraries. The “no DNA” condition is also referred as the “second isolation” used in the fitness analysis.

### DNA library preparation and sequencing

Libraries were prepared as previously described (79). Mutant libraries from the −80°C frozen stock were thawed and centrifuged at maximum speed to pellet them. gDNA extraction was carried out directly on the pellet cells with the Wizard Genomic DNA Purification Kit (Promega) according to manufacturer’s instructions. ∼30 µg of DNA were mechanically sheared by sonication using a Branson sonifier for 4 min (1 sec on and 11 sec off; 20 % intensity) in 0,5-mL PCR tubes kept on ice. Small gDNA molecules were removed by mixing sonicated gDNA with 0.6X Agencourt Ampure XL magnetic beads (Beckman Coulter) according to manufacturer’s instructions. These treatments led to gDNA fragments being between 300 and 1000 pb. Homopolymeric cytosine-tails (C-tail) were then added to the 3’ ends of all fragments by incubation of 3 µg of size-selected DNA fragments with the recombinant terminal deoxynucleotidyl transferase (rTdT, 30 U/µL, Promega) at 37°C for 1H, followed by heat inactivation at 75°C for 20 min. TdT reagents were then removed by purifying the TdT reaction mixture with 1X of Ampure XL beads. To amplify transposon junctions, a first-round of PCR (PCR1) was performed in a final volume of 50 µL by mixing 500 ng C-tailed DNA, 1 μL biotinylated pBT20-PCR1 primer (30 μM), 3 μL olj376 primer (30 μM), 2.5 μL dNTPs (10 mM), 10 μL Q5 reaction buffer and 0.75 μL Q5 High-Fidelity DNA Polymerase (New England Biolabs). PCR1 products were purified using 1X Ampure beads. Biotinylated and purified PCR1 products were then selectively captured using Dynabeads M-280 Streptavidin (Invitrogen) according to manufacturer’s instructions. A second-round of PCR was carried out in a final volume of 50 µL by resuspending Dynabeads (which have PCR1 products bound to them) in the pre-prepared PCR2 reaction mix constituted of 1 μL pBT20-PCR2 primer (30 μM), 1μL TdT_index_X primer (30 μM), 2.5 μL dNTPs (10 mM), 10 μL Q5 reaction buffer and 0.75 μL Q5 High-Fidelity DNA Polymerase. PCR2 products were purified with 1X Ampure XL beads. The obtained libraries were sequenced on an Illumina HiSeq 4000 in single-end 50 pb using the custom sequencing primer Read1TnLp. Samples and conditions are listed in Dataset S1. Essentiality analysis was performed using reads from sample XRCR13. Fitness analysis was performed by comparing reads from samples XRCR24, 26, 36 and 38 (no DNA conditions from the transformation screen) versus XRCR13. Analysis of transformation was performed by comparing samples XRCR27, 39 (legK2::kan transforming DNA) vs samples XRCR26, 38 (no DNA control) and XRCR25, 37 (ihfB::kan transforming DNA) to samples XRCR24, 36 (no DNA control). Raw sequencing reads were deposited to the European Nucleotide Archive (https://www.ebi.ac.uk/ena/) under the study accession number PRJEB40244.

### Tn-seq data analysis

For each condition, 10-50 million reads were obtained and trimmed with tools from the Galaxy’s project public server. Fastx_clipper was used to cut poly-C tails and remove short reads (<15 pb after polyC clipping). Then, reads were filtered by quality using trimmomatic and quality checked with FastQC. Trimmed reads in the fastq output file were mapped to the reference genome using Tn-seq software TPP (Tn-seq pre-processor) (80). Output wig files from TPP were used to perform essentiality analysis using Transit (81). Single-condition essentiality analysis was performed with the hmm (46) or Gumbel (45) methods. Conditional essentiality analysis was performed with the “resampling” method according to the Transit software documentation. Complete genome sequence of HL-0709-3014 was obtained (see *Genome sequence and accession numbers*) and annotated with Prokka (82). An orthology search was carried out between the strains of *L. pneumophila* HL77S, Paris and Philadelphia-1 using the orthology detection eggNOG mapper (83) and COG and KEGG number were assigned to each gene.

### Transformation assays

Natural transformation assays were conducted differently depending on the genetic background of *L. pneumophila strain* used: (1) For the constitutively transformable *rocC*_TAA_ strains, natural transformation was conducted on solid medium at 37°C as follows. The strains were streaked on CYE solid medium from a frozen stock culture and incubated for 72 h at 37°C. The strains were then restreaked on a new CYE plate and incubated overnight at 37°C to obtain freshly growing cells. Bacteria were resuspended in sterile water to an OD600 of 1 to obtain a suspension of 1.10^9^ colony forming units (CFUs) per milliliter. 10 µL of the suspensions (∼1.10^7^ CFU) were spotted on CYE with 1.5 µg of transforming DNA. Once the spots are absorbed by the agar, plates were incubated at 37°C for 24 h. Each spot was resuspended in 200 µL sterile water and used to perform tenfold serial dilutions which were then plated on nonselective medium and selective medium. Plates were incubated at 37°C for 72 h. Finally, transformation frequencies were calculated as the ratio of the number of CFUs counted on selective medium divided by the number of CFUs counted on nonselective medium. For all the *rocC*_TAA_ strains, “rpsL” PCR product was used as transforming DNA. This transforming DNA is obtained by amplificating the 2-kB regions upstream and downstream the *rpsL* single point mutation conferring resistance to streptomycin (PCR primers pairs rpsL_F/rpsL_R). Transformation experiments on strains bearing the p0681F and p1890F plasmids were performed the same way, using CYE plates containing different concentration of IPTG. (2) For the non-constitutively transformable strains of *L. pneumophila*, transformation was realized in liquid medium at 30°C as follows: strains were streaked on CYE solid medium from a frozen stock culture and incubated for 72 h at 37°C and then restreaked on a new CYE plate and incubated overnight at 37°C. Fresh bacteria were resuspended in 3 mL of AYE in 13-mL tubes to an OD∼0.2 with 2 µg of transforming DNA and cultivated at 30°C with shaking for 24 h. Tenfold serial dilution of each culture was then performed and plated on nonselective medium and selective medium and incubated at 37°C for 72 h. Finally, transformation frequencies were determined as described above. (3) For *letA* mutants of constitutively and non-constitutively transformable strains of *L. pneumophila*: strains were streaked on CYE solid medium from a frozen stock culture and incubated for 72 h at 37°C and then restreaked on a new CYE plate and incubated overnight at 37°C. Fresh bacteria were resuspended in 3 mL of AYE in 13-mL tubes to an OD∼0.2 and cultivated at 30°C with shaking until OD∼2-4 (corresponding to the competence phase of *L. pneumophila*). A volume corresponding to 1.10^8^ bacteria was spotted on CYE plates with 1.5 µg the *rpsL* PCR product. The following steps were the same as for the transformation of constitutively transformable *rocC*TAA strains as mentioned in (1).

### Detection of extracellular pilin by Western-Blot

Strains bearing plasmid p1890F were grown overnight at 37°C on CYE containing 500 µM IPTG, and were then resuspended in 2 mL AYE at an OD600∼1.5. 1 mL of the suspension was then submitted to max-speed vortex agitation for 1 min, and centrifuged 15 min at 21,000 g and 4°C. Supernatants were recovered in a new tube and centrifuged again, while pellets were saved on ice. After centrifugation, 900 µL of supernatants were recovered and proteins were precipitated by adding 100 µL of Trichloroacetic Acid (TCA, final concentration of 10%). After 30 min of incubation on ice, a 15 min centrifugation at 21,000 g and 4°C was performed. Pellets were washed three times with acetone, dried at room temperature and resuspended with 100 µL of Laemmli Sample Buffer 1X. Pellets previously saved on ice were resuspended with 150 µL of Sample Buffer 1X. Samples were then analyzed by Western-blot. Aliquots were boiled for 5 min and subjected to denaturing polyacrylamide gel electrophoresis. Proteins from SDS-polyacrylamide gels were electrophoretically transferred to nitrocellulose membranes (Schleicher and Schuell) and subsequently stained with Ponceau S (Sigma) to check the loading of the lanes. Membranes were incubated with monoclonal Anti-FLAG antibody (dilution 1:1000, Sigma F1804) as a primary antibody and an anti-mouse peroxidase conjugate (dilution 1:50000, Sigma A0168) as secondary antibody. Nitrocellulose membranes were revealed with the SuperSignal® West Dura detection system (Pierce) and an imaging workstation equipped with a charge-coupled device camera (Thermo).

### Determination of the DNA uptake ability

The ability of the transformation-deficient mutants to uptake DNA was determined as follows: strains were inoculated in AYE media at an OD600 = 0.05 and tubes were incubated overnight under constant shaking at 30°C. When OD600 = 0.9 was reached, 1 mL of each culture was centrifuged 3 min at 5000g, and pellets were resuspended in 200 µL ultrapure water containing 2 µg of pGEM-HYG1. This plasmid is non-replicative plasmid in *L. pneumophila* and, as it contains no homology with *L. pneumophila* genome, it cannot integrate by recombination either.. After 20 min of incubation at 37°C, tubes were centrifuged 3 min at 5000g and pellets were resuspended in 200 µL AYE liquid medium containing 10 Units of DNase I (Sigma). After 20 min of incubation at 37°C, DNase I was removed and bacteria were washed by two successive centrifugation 3 min at 5000 g and resuspension in 1 mL of water. Pellets were finally resuspended in 100 µL ultrapure water, and incubate 30 min at 65°C to complete DNAse I inactivation and kill bacteria. DNA uptake ability of each mutants was then determined by PCR, using two couples of primers amplifying on the one hand the chromosomal *mreB* gene (*lpp0873*) and on the other hand a part of pGEM-HYG1, giving respectively PCR products of 1194 pb (mreBseqF/mreBseqR) and 1657 pb (M13F(−47)/M13R(−48)).

### Microscopy

Bacteria expressing the FLAG-tagged pilins were grown as spots on CYE plates with 0.5 mM IPTG for 24 h at 37°C. Bacteria were gently collected with a pipette tip. In order to limit shearing and breaking of the pilus, the pipette tip was left standing an eppendorf tube with 1 mL of water for a few minutes. Once the collected bacterial culture is starting to dissociate and falling off from the tip, the bacterial pellet is resuspended gently by slowly pipetting up and down. The collected 1 mL suspensions were centrifuged 3 min at 5000 g and pellets were gently resuspended in 300 µL PBS Formaldehyde 3.7% and incubated at room temperature for 30 minutes. Acid-washed (ethanol/HCl 1M) glass coverslips were coated with poly-L-lysine by immersion in a poly-L-lysine 0.01% solution in distilled water (Sigma-Aldrich) for 5 minutes. Fixed bacteria in PBS Formaldehyde 3.7% were pipetted (250 µl) on the air-dried coverslips and let to settle and stick to the coverslips for about 30 minutes. Coverslips were then washed twice with PBS, and incubated with monoclonal anti-FLAG M2 fluorescein conjugates at 1/200 in PBS for 1h. Coverslips were then washed twice with PBS and DNA was labeled using Hoechst 33288 (12 µg/mL in PBS) 1 h. Coverslips were washed twice in PBS and mounted using 8 µL of mounting solution (DAPCO). After an overnight incubation at 4°C slides were observed and imaged with an epifluorescence microscope (Zeiss Axioplan 2).

### Genome sequence and raw reads accession numbers

The complete genome of isolate HL-0709-3014 was obtained using Illumina MiSeq paired end reads from previously available SRA sample ERS1305867 and long reads from Oxford Nanopore sequencing on a MinION sequencer according to manufacturer’s instructions (Oxford Nanopore). Illumina and Nanopore reads were then used for short reads/long reads hybrid assembly using Unicycler v0.4.6 (84). The complete genome of isolate HL-0709-3014 is available under accession numbers CP048618.1 (chromosome) and CP048619.1 (plasmid). The strain is listed under the name *Legionella pneumophila* strain ERS1305867 (BioProject: PRJEB15241; BioSample: SAMEA4394418). Raw sequencing reads of the Tn-seq samples are available at the European Nucleotide Archive (https://www.ebi.ac.uk/ena/) under the study accession number PRJEB40244.

## Acknowledgements

We warmly thank Christophe Ginevra (Centre National de Référence des Légionelles) for kindly providing Nanopore sequencing reads and full genome assembly of HL-0709-3014. We also thank Chloé Vallantin for technical assistance with clinical isolates and Annelise Chapalain and Johann Guillemot for providing mouse and human monocytes cultures and helpful advice on infection with *L. pneumophila*. We thank Vladimir Shevchik for his insight into type IV secretion and minor pills. We acknowledge Laetitia Attaiech and Maria-Halima Laaberki for their critical assessment of the manuscript. This work was supported by the LABEX ECOFECT (ANR-11-LABX-0048) of Université de Lyon, within the program “Investissements d’Avenir” (ANR-11-IDEX-0007) operated by the French National Research Agency (ANR).

LH, PAJ and BCG designed and performed experiments, and analyzed data. LH and XC analyzed Tn-seq data. LH and XC wrote the manuscript. XC conceptualized and supervised the project.

---

**Figure S1.**
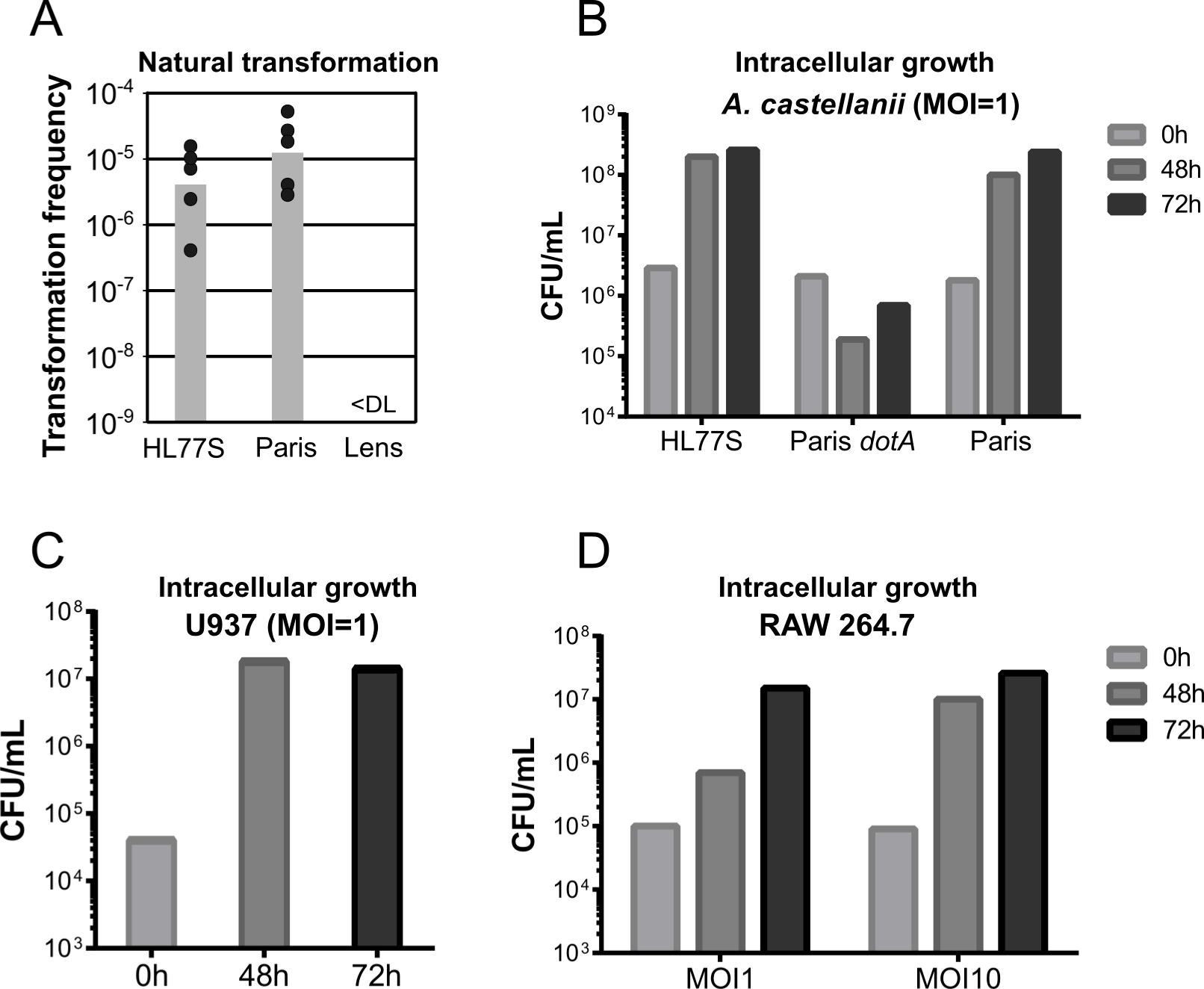
Phenotypic characterization of strain HL77S, a streptomycin-resistant mutant of the clinical isolate HL-0709-3014. A) Natural transformability of HL77S compared to the Paris and Lens strain. Transformation was tested by growing the strains in AYE at 30°C for 24h in the presence of 2 µg of transformation DNA consisting of a kanamycin resistance gene interrupting the *ihfB* gene. B) Intracellular replication of HL77S in the amoeba *Acanthamoeba castellanii*. Cells were infected with a suspension of HL77S at a multiplicity of infection (MOI) of 1. At the initial time point, after 48 h and 72 h of culture at 30°C, colony-forming units are determined by plating on CYE medium. C) Intracellular replication of HL77S in differentiated human monocytes of the U937 cell line. Cells were infected with a suspension of HL77S at a MOI of 1. At the initial time point, after 48 h and 72 h of culture at 30°C, colony-forming units are determined by plating on CYE. D) Intracellular replication of HL77S in the murine macrophage-like cell line RAW 264.7. Cells were infected with a suspension of HL77S at a MOI of 1 or 10. At the initial time point, after 48 h and 72 h of culture at 30°C, colony-forming units are determined by plating on CYE.

**Table S1.**
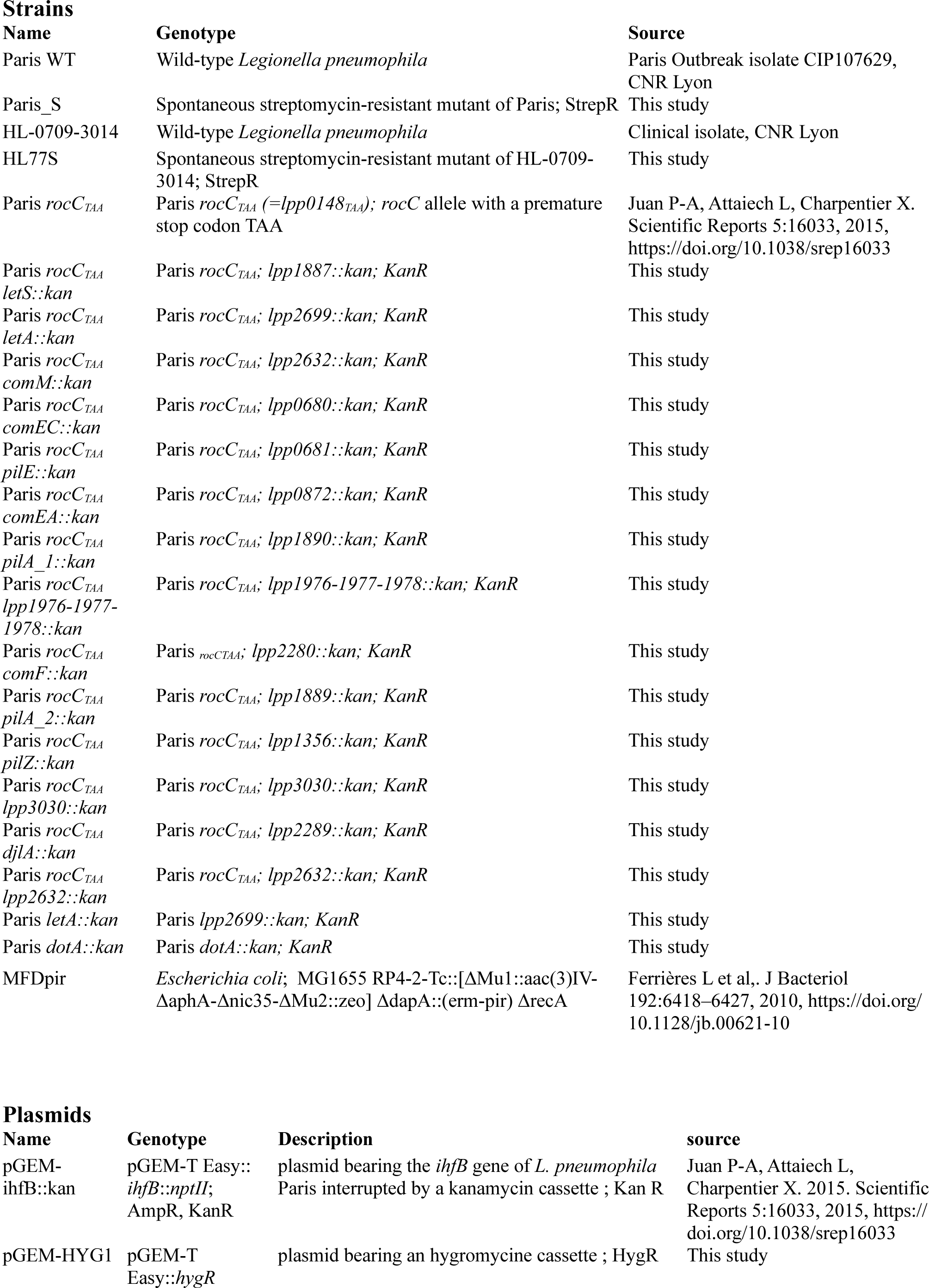
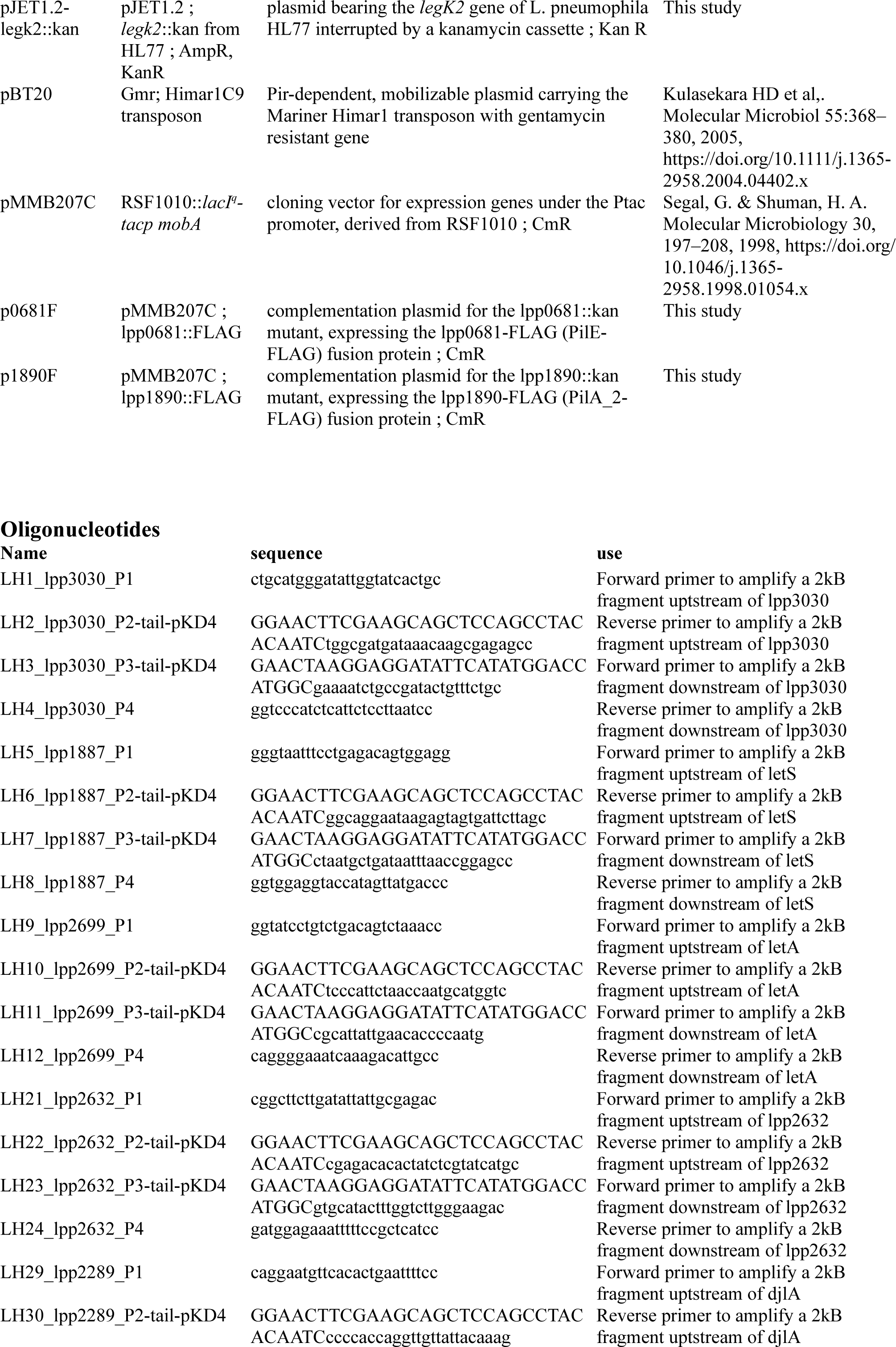
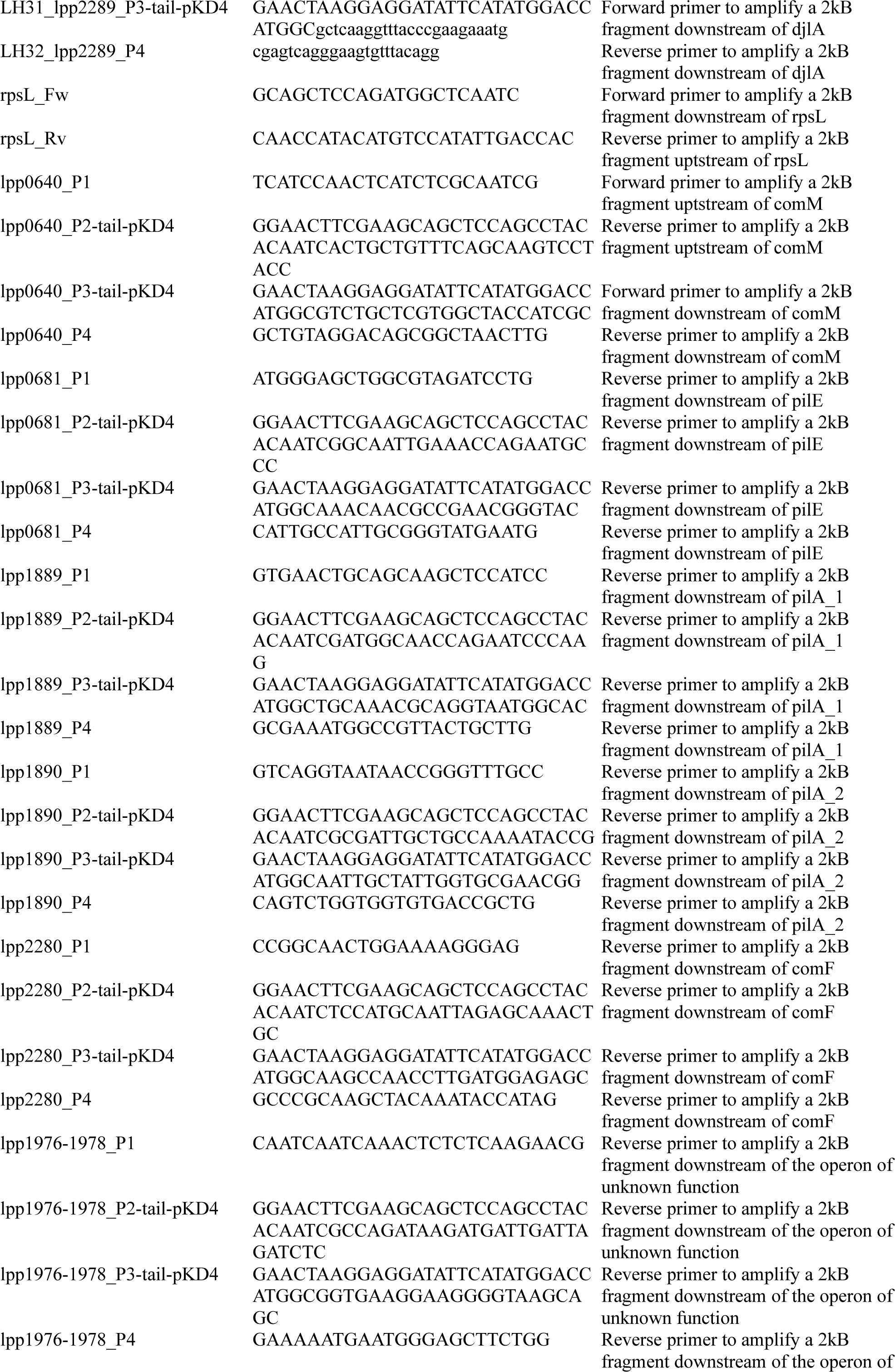
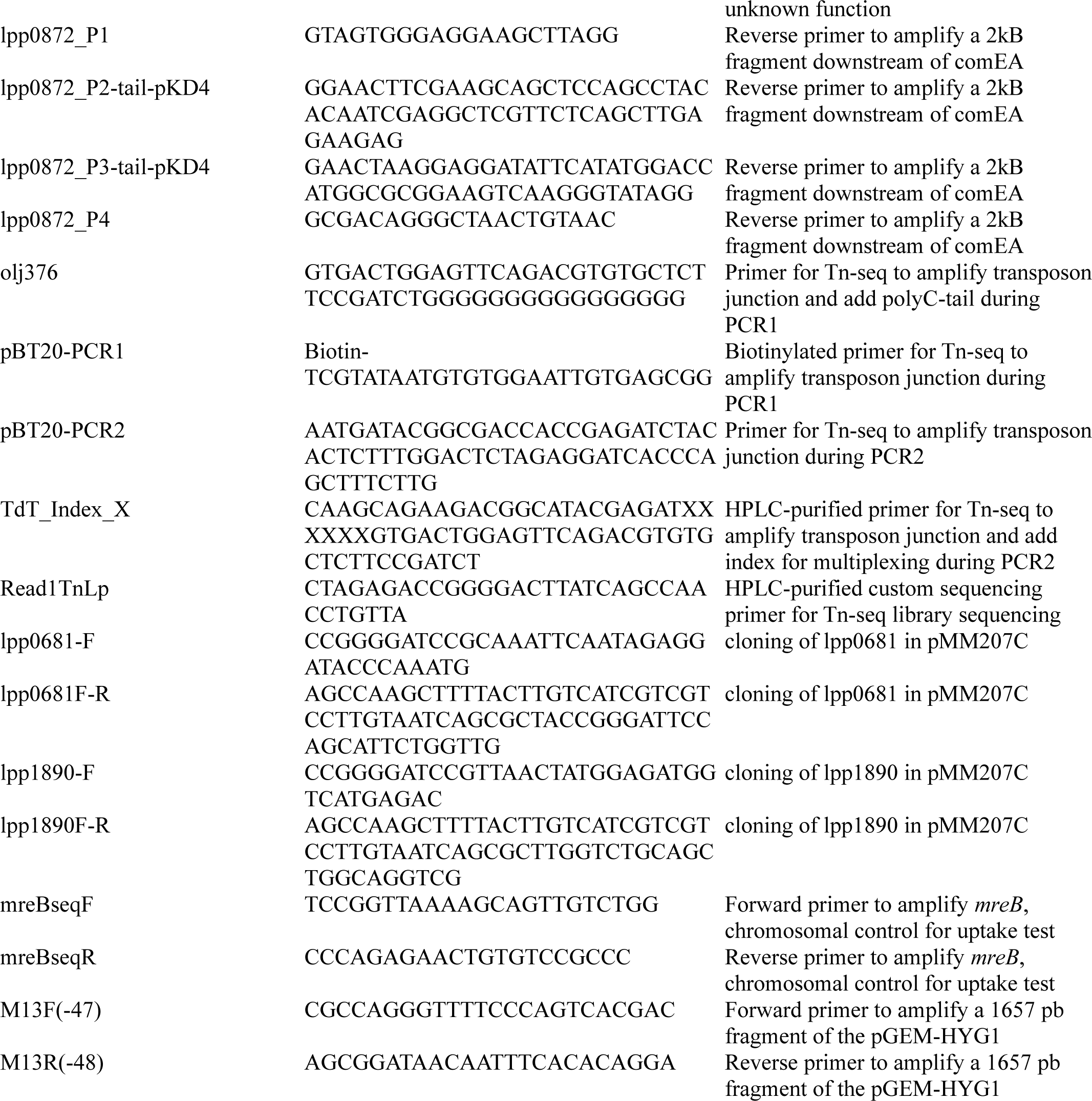
Strains, plasmids and oligonucleotides used in this study.

